# Knockdown of Hypocretin/Orexin Attenuates Extended-Access Cocaine Self-Administration in Rats

**DOI:** 10.1101/184911

**Authors:** Brooke E. Schmeichel, Alessandra Matzeu, Pascale Koebel, Leandro F. Vendruscolo, Brigitte L. Kieffer, George F. Koob, Rémi Martin-Fardon, Candice Contet

## Abstract

The hypocretin/orexin (HCRT) neuropeptide system regulates feeding, arousal state, stress responses, and reward, especially under conditions of enhanced motivational relevance. In particular, HCRT neurotransmission facilitates drug-seeking behavior in circumstances that demand increased effort and/or motivation to take the drug. The present study used a shRNA-encoding adeno-associated viral vector to knockdown *Hcrt* expression throughout the dorsal hypothalamus in adult rats and determine the role of HCRT in cocaine self-administration. Longterm *Hcrt* silencing did not impact cocaine self-administration under short-access conditions, but robustly attenuated cocaine intake during extended self-administration access, a model that mimics key features of compulsive cocaine-taking. In addition, *Hcrt* silencing decreased motivation for both cocaine and palatable food (i.e., sweetened condensed milk; SCM) under a progressive ratio schedule of reinforcement, but did not alter responding for SCM under a fixed ratio schedule. Importantly, *Hcrt* silencing did not affect food or water consumption, and had no consequence to general measures of arousal-dependent behaviors. **At the molecular level, longterm *Hcrt* knockdown moderately reduced the downstream expression of dynorphin (DYN) and melanin-concentrating hormone (MCH) in the dorsal hypothalamus.** These original findings support the hypothesis that HCRT neurotransmission promotes operant responding for both drug and non-drug rewards, preferentially under conditions requiring a high degree of motivation. Furthermore, the current study provides compelling evidence for the involvement of the HCRT system in cocaine self-administration also under low-effort conditions in rats allowed extended access, **possibly via functional interactions with DYN and MCH signaling**.

## Introduction

Independently discovered in 1998 by de Lecea and Sakurai, hypocretin/orexin peptides hypocretin-1 (HCRT-1, also ORX-A) and hypocretin-2 (HCRT-2, also ORX-B), are derived from a common precursor, prepro-HCRT. These HCRT neuropeptides are synthesized in well-defined subregions of the dorsal hypothalamus: lateral hypothalamus proper, dorsomedial hypothalamus, and perifornical area (de Lecea *et al*, 1998; Peyron *et al*, 1998; Sakurai *et al*, 1998). HCRT projections are found throughout the brain and in regions known for their involvement in arousal, stress, and drug and non-drug reinforcement. These areas include, but are not limited to, the central amygdala, nucleus accumbens, ventral tegmental area, arcuate nucleus and paraventricular nucleus of the hypothalamus (Baldo *et al*, 2003; Nambu *et al*, 1999; Peyron *et al*, 1998; Schmitt *et al*, 2012). HCRT neuropeptides bind two G-protein-coupled receptors, HCRT-receptor 1 and -receptor 2 (HCRT-R1 and HCRT-R2, respectively; Sakurai *et al*, 1998) that also are distributed widely throughout the brain (Marcus *et al*, 2001; Trivedi *et al*, 1998). Accordingly, the HCRT system is involved in a multitude of physiological functions, such as the regulation of feeding, arousal, sleep/wake states, stress responses, energy homeostasis, and reward (for review, see Berridge and España, 2005; Boutrel and de, 2008; Johnson *et al*, 2010; Tsujino and Sakurai, 2009).

An abundant body of literature demonstrates the critical importance of HCRT transmission in the consumption and seeking of various reinforcers, including cocaine (Boutrel *et al*, 2005; España *et al*, 2010; Hollander *et al*, 2012; Martin-Fardon and Weiss, 2014a; Muschamp *et al*, 2014; Prince *et al*, 2015; Smith *et al*, 2009, 2010; Wang *et al*, 2009; Zhou *et al*, 2012; Martin-Fardon *et al*, 2016; Schmeichel *et al*, 2016), nicotine (Hollander *et al*, 2008; LeSage *et al*, 2010; Plaza-Zabala *et al*, 2010, 2013), alcohol (Anderson *et al*, 2014; Brown *et al*, 2013; Dhaher *et al*, 2010; Jupp *et al*, 2011; Lopez *et al*, 2016; Martin-Fardon and Weiss, 2014b; Olney *et al*, 2015), heroin (Schmeichel *et al*, 2015; Smith and Aston-Jones, 2012), sucrose, and saccharin (Cason and Aston-Jones, 2013a, 2013b; Olney *et al*, 2015). Importantly, HCRT-receptor blockade generally does not influence drug self-administration under continuous, low-effort reinforcement, but rather blocks self-administration when the contingency of reinforcement requires higher levels of motivation to acquire the drug (Mahler *et al*, 2014). However, recent studies have indicated that acute blockade of HCRT signaling reduces not only the appetitive aspect but also the consummatory aspect of drug taking in dependent animals (Lopez *et al*, 2016; Schmeichel *et al*, 2015, 2016).

The aim of the present study was to investigate the role of HCRT neurotransmission in cocaine self-administration when rats are given extended-access to the drug. The extended-access model produces a gradual escalation of cocaine self-administration and an increased motivation to obtain cocaine, along with increases in brain self-stimulation thresholds during withdrawal, stress reactivity, resistance to punishment and reinstatement susceptibility (Ahmed *et al*, 2002; Ahmed and Koob, 1998; Aujla *et al*, 2008; Jonkman *et al*, 2012; Mantsch *et al*, 2004, 2008; Orio *et al*, 2009; Pelloux *et al*, 2007; Vanderschuren and Everitt, 2004; Wee *et al*, 2008). To examine the role of HCRT transmission in this model, *Hcrt* expression was silenced longterm throughout the dorsal hypothalamus of adult rats using a short hairpin RNA (shRNA)-encoding adeno-associated viral (AAV) vector. To further investigate the function of HCRT in reward consumption and potentially dissociate its role in regulating motivation for drug versus food, the effects of *Hcrt* silencing on self-administration of a highly palatable food reinforcer (sweetened condensed milk, SCM), as well as for regular food pellets and water, were assessed. In addition, given the modulatory role of HCRT in a multitude of behavioral and physiological functions, locomotor activity, anxiety-like behavior, and stress-induced analgesia and corticosterone response were measured to further evaluate the specificity of the behavioral consequences of *Hcrt* knockdown. **Finally, molecular adaptations to prolonged reduction in HCRT signaling were investigated by analyzing the expression of prodynorphin (PDYN) and melanin-concentrating hormone (MCH), two neuropeptides also synthesized in the dorsal hypothalamus.**

## Material and Methods

### Animals

Forty adult male Wistar rats (Charles River, Raleigh, NC), weighing between 225-275 g at the beginning of the experiments, were housed in groups of 2-3 per cage in a temperature-controlled (22ºC) vivarium on a 12 h/12 h light/dark cycle (lights on at 18:00 h) with *ad libitum* access to food and water. The rats acclimated to the animal facility for at least 7 days before surgery. All procedures adhered to the National Institutes of Health Guide for the Care and Use of Laboratory Animals and were approved by the Institutional Animal Care and Use Committee of The Scripps Research Institute.

### Viral Vectors

Recombinant shRNA-encoding AAV vectors were produced using an AAV helper-free system (Stratagene, France; as described in Darcq *et al*, 2011). In these vectors, the shRNA sequence is expressed under the control of the mU6 promoter, while the enhanced green fluorescent protein (GFP) is expressed under the control of the cytomegalovirus promoter to label transduced cells. The shRNA sequence targeting the *Hcrt* transcript (shHCRT; 5’-GTCTTCTATCCCTGTCCTAGT-3’) was selected using the BLOCK-iT RNAi Designer algorithm (ThermoFisher). A scrambled sequence (shSCR; 5’-GCTTACTTTCGGCTCTCTACT-3’) was used as negative control. Loop sequence was 5’-AGTCGACA-3’ for both. **An AAV2 serotype (titer of 7.4 x 10**^**11**^**GU/mL**) was used to characterize the time-course of *Hcrt* knockdown, while AAV5 vectors (titers of 2.4 x 10^12^ for shHCRT and 2.6 x 10^12^ GU/mL for shSCR) were used for all behavioral experiments.

### AAV Injections

Rats were anesthetized with isoflurane (1-3%), mounted in a stereotaxic frame (Kopf Instruments, Tujiunga, CA). A stainless steel 30-gauge double injector was used to inject viral vectors at two mediolateral levels (AP -2.9 mm and ML ± 0.5/1.75 mm from bregma, and DV - 8.7 mm from dura; Paxinos and Watson, 2013). Bilateral injections of AAV5-shSCR or AAV5-shHCRT were performed for all behavioral experiments. **Unilateral injections of AAV2-shHCRT were performed for *Hcrt* knockdown time-course characterization, with brains collected 2 weeks (*n* = 3), 4 weeks (*n* = 2), or 6 weeks (*n* = 3) after AAV2 injection (see Supplementary Figure S1)**. Injections were made using a micro-infusion pump (Harvard 22 Syringe Pump, Holliston, MA) with a flow rate of 0.5 μl/min over 4 min (2 μl/site). Injectors remained in place for 10 min to assure adequate diffusion of the solution and prevent backflow along the injector track.

### Experimental Design

Three distinct cohorts were used to test the effect of long-term *Hcrt* silencing on cocaine self-administration under short-and long-access schedules (*n* = 8 each shSCR and shHCRT, three rats were excluded from each group due to misplaced injections and/or catheter patency failures), SCM self-administration (*n* = 6 each shSCR and shHCRT, one rat was excluded from each group due to misplaced injections), and general behavior (*n* = 6 each shSCR and shHCRT, one rat was excluded from each group due to misplaced injections. General behavioral testing included food/water self-administration, locomotor activity, anxiety-like behavior, and stress-induced analgesia and corticosterone response). **The timeline of all behavioral testing in each cohort is shown in Supplementary Figure S2.** Behavioral testing resumed 2-3 weeks after AAV5 injection and was conducted during the dark phase of the circadian cycle, unless otherwise noted.

### Cocaine Self-administration

Rats were surgically prepared with indwelling jugular vein catheters (Dow Corning, Midland, MI) and intravenous self-administration sessions were conducted as previously described (Wee *et al*, 2012; Zorrilla *et al*, 2012; see Supplementary Methods). Briefly, rats were trained to press one of the two levers on a fixed ratio 1 (FR1) schedule of reinforcement to obtain 0.1 ml of cocaine (0.50 mg/kg/infusion) per response in 1 h sessions. After the acquisition of cocaine self-administration, rats were injected with AAV vectors and allowed to recover for approximately 2 weeks (see Figure S1A). Rats were then given short access (ShA; 1 h) to cocaine self-administration for 11 sessions and then transitioned to long access (LgA; 6 h) to cocaine for an additional 14 sessions (Figure S2A). Testing under a progressive ratio (PR; see Supplementary Methods) schedule of reinforcement occurred following LgA sessions.

### Sweetened Condensed Milk (SCM) Self-administration

SCM (Nestlé USA, Inc., Solon, OH) self-administration training occurred in daily 30-min sessions on a FR1 TO20 schedule of reinforcement, prior to AAV5 injection. Sessions were initiated by the extension of both levers into the operant chamber, and responses on the active lever resulted in the delivery of SCM (0.1 ml; 2:1 v/v in water) into a drinking receptacle. Responses on the inactive lever were recorded but had no scheduled consequences. Following AAV injection, the rats were allowed to recover for 3 weeks after which SCM self-administration resumed in 24 daily 30-min sessions (see Figure S2B). The rats were then tested on a PR schedule of reinforcement using the same ratio described for cocaine.

### Food/water Self-administration

Rats underwent three food/water self-administration sessions (22 h/day, 11 h dark/11 h light), preceded by two days of habituation. Operant boxes (22 cm x 22 cm x 35 cm) were equipped with two holes. Rats were allowed to make nose poke responses in order to obtain food pellets (MLab Rodent Tablet 45 mg, TestDiet) from the pellet dispenser (hole on right wall; FR3) or 0.1 ml water from the water dispenser (hole on left wall; FR1). Responses were detected by photobeams mounted in the holes and recorded automatically. Between sessions, animals stayed in their home cage. Food/water self-administration was examined before and after AAV5 injection (Figure S2C).

### Locomotor Activity

Locomotor activity was tested in photocell-equipped wire mesh cages holding two photobeams along the lateral walls. Locomotor activity was recorded for three consecutive days (22 h/day). Food and water were available *ad libitum*. Computer-recorded photobeam breaks were analyzed as a measure of locomotor activity. Crossovers were defined as two consecutive photobeam breaks as the rat moved from front to rear of cage, or vice versa. Detailed descriptions of procedures are provided in the Supplementary Methods. Locomotor activity was monitored before and after AAV5 injection (Figure S2C).

### Elevated Plus Maze

The elevated plus maze apparatus (Kinder Scientific, Poway, CA) comprised four arms (two closed and two open arms). At the beginning of the test, each rat was placed in the center of the maze facing a closed arm and behavior was video recorded for 5 min. Between rats, the apparatus was cleaned with water and dried. Detailed descriptions of procedures are provided in the Supplementary Methods. Time (seconds) spent in each arm was recorded. Elevated plus maze was tested only after AVV5 injection (Figure S2C).

### Stress-induced Analgesia Testing

Stress-induced analgesia testing was conducted as previously described (Vendruscolo *et al*, 2004; detailed descriptions of procedures are provided in the Supplementary Methods). Briefly, each rat was placed individually on a hot plate set to 54°C. Time (s) to hind-paw lick was recorded, with a cut-off time of 60 s. One hot plate test was conducted before the forced swim test (PRE) and one test 10 min following forced swim (POST). For forced swim, rats were individually placed in a Plexiglas cylinder filled with water for 5 min. Water was replaced between subjects. Rats underwent stress-induced analgesia testing only after AVV5 injection (Figure S2C).

### Plasma corticosterone

Two hours following the elevated plus maze testing (pre-swim condition) and 20 min after the forced swim test (post-swim condition) blood samples were collected for corticosterone measurements (Figure S2C). In brief, rats were restrained and tail blood (approximately 0.2 ml) was collected into tubes coated with a 10% ethylenediaminetetraacetic acid solution and centrifuged immediately. Plasma was isolated and stored at -80°C. Plasma corticosterone concentrations were determined using a Corticosterone Enzyme Assay Kit (Arbor Assays, Ann Arbor, MI) according to the manufacturer's protocol.

### Immunohistochemistry

Detailed descriptions of immunohistochemical procedures and cell counting are provided in the Supplementary Methods. Briefly, rats were transcardially perfused with paraformaldehyde, brains were post-fixed, and six series of 40 μm coronal sections were collected in vials. For each behavioral cohort, two series were processed for immunohistochemistry using antibodies directed against GFP (Abcam; 1:10,000) and preproHCRT (EMD Millipore; 1:1000). In the cohort used for general behavior testing, a third series was used to analyze MCH immunoreactivity (Santa Cruz Biotechnology; 1:1000). GFP immunoreactivity was used to evaluate the location and extent of viral transduction in each rat at the end of the experiment **and was consistently identified throughout the HCRT neuronal field (dorsal hypothalamus)**. HCRT-positive and MCH-positive cells were counted manually at 20x magnification.

### *In situ* hybridization

Brains were snap-frozen in isopentane and ten series of 20 μm coronal cryostat sections were collected on Superfrost Plus slides. Three series were processed for *in situ* hybridization with probes directed against GFP, preproHCRT (*Hcrt*), and prodynorphin (*Pdyn*) mRNAs. Plasmids containing the rat *Hcrt* and *Pdyn* cDNAs were kindly donated by Dr. Joel Elmquist at UT Southwestern (Elias *et al*, 2001; Sakurai *et al*, 1998). Digoxigenin (DIG)-labeled riboprobes were synthesized using a kit (Roche, Indianapolis, IN). Chromogenic *in situ* hybridization was conducted as described in Herman *et al*, 2015. The GFP signal was used to define the boundaries of viral transduction in each brain section. Numbers of *Hcrt*-positive and *Pdyn*-positive cells in adjacent sections were then counted within the transduced area using Stereo Investigator (MBF Bioscience) at 20x magnification.

### Statistical Analysis

Statistical analyses were performed using Prism 7 (GraphPad Software, La Jolla, CA). All data are expressed as means and standard errors of the mean (+SEM). **Cell counts from the knockdown time-course experiment were analyzed by two-way ANOVA, with viral vector (shHCRT-injected versus non-injected control side) as within-subjects factor and time (2, 4 and 6 weeks) as between-subjects factor.** Fixed ratio self-administration, locomotor activity, hot plate, and corticosterone data were analyzed using a repeated-measures two-way analysis of variance (ANOVA), with viral vector treatment (shHCRT and shSCR) as the between-subjects factor and time as the within-subjects factor. When appropriate, *post hoc* comparisons were performed using Sidak's multiple-comparison test. Cell counts in behavioral cohorts and elevated plus maze data were analyzed using an unpaired, two-tailed Student's *t* test. Progressive ratio self-administration data were analyzed using a Mann-Whitney U Test. P < 0.05 was considered statistically significant for all tests.

## Results

### Characterization of *Hcrt* knockdown

The AAV5 transduction spread and knockdown efficiency were verified in each rat at the end of each behavioral experiment. GFP immunoreactivity was consistently identified throughout the HCRT neuronal field (Figure 1A). Bilateral injections of AAV5-shHCRT robustly silenced HCRT expression (Figure 1B, C). The number of HCRT cells was reduced by more than 80% compared to shSCR rats in each behavioral cohort (Figure 1D, *t*_(8)_ = 6.42, p < 0.001; Figure 1E, *t*_(8)_ = 5.10, p < 0.001; Figure 1F, *t*_(8)_ = 4.13, p < 0.01).

**Figure 1.**
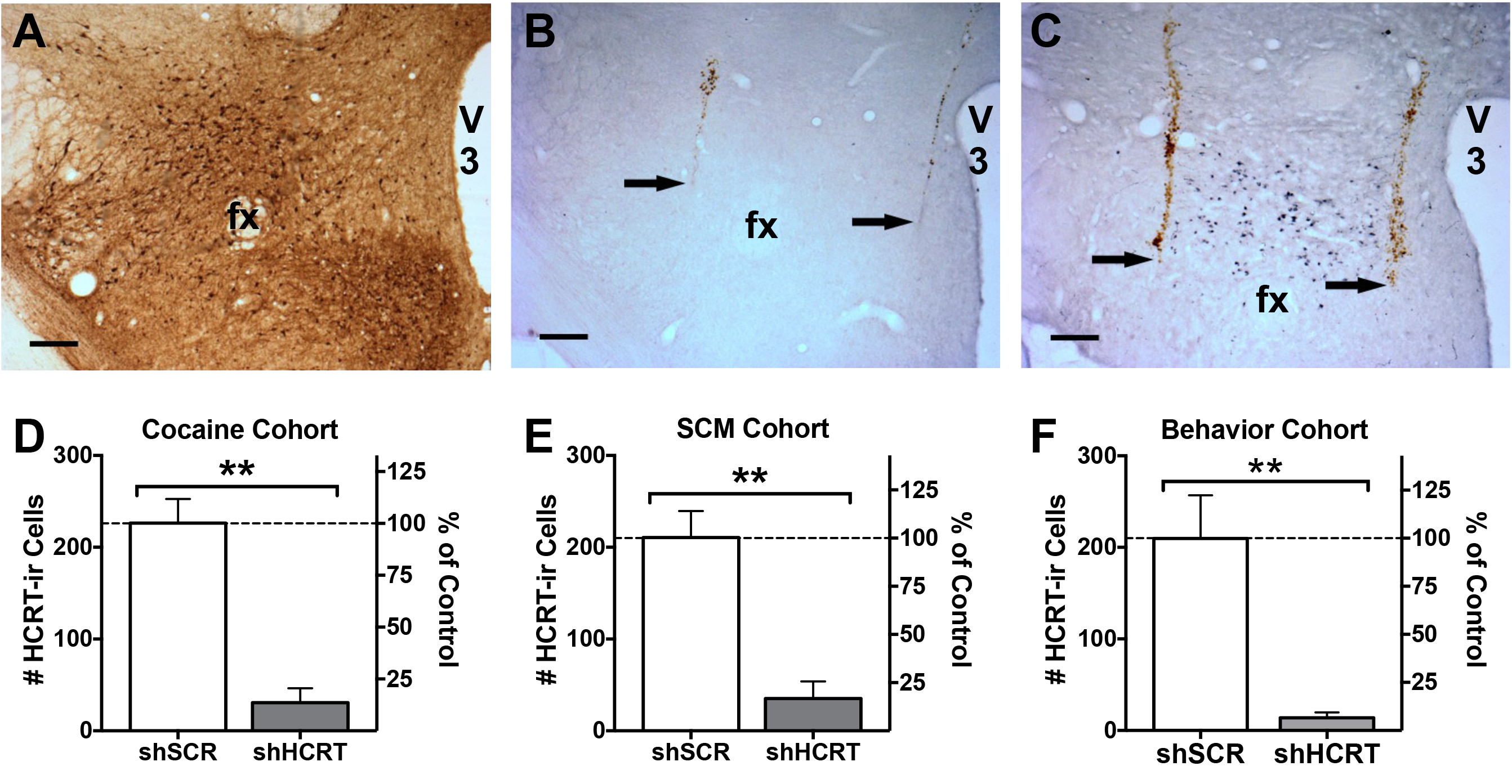
Histological characterization of viral transduction and quantification of *Hcrt* knockdown. Rats received double bilateral injections of AAV5-shSCR or AAV5-shHCRT (*n* = 5, each) and were then subjected to behavioral testing. Brains were collected at the end of each experiment for verification of viral transduction by GFP immunohistochemistry (A, brown precipitate). The area of GFP expression consistently encompassed the entire hypothalamus, including areas known to contain HCRT neurons (i.e., lateral hypothalamus proper, perifornical area, and dorsomedial hypothalamus). Adjacent sections were processed for prepro-HCRT immunolabeling (B-C, blue-gray precipitate) to quantify HCRT knockdown in each rat. (A-C) Images were captured at a 5x magnification, scale bar = 250 μm. Ventral-most end of injector track is indicated with arrows. fx, fornix; V3, third ventricle. (D-F) shHCRT reduced the number of HCRT neurons by more than 80% in all three cohorts. Bars represent the mean (+SEM) number of preproHCRT-immunoreactive (HCRT-ir) neurons (left axis) and the percentage of the control group (shSCR; right axis) across three coronal sections. **p<0.01 versus shSCR.

### *Hcrt* knockdown reduces low-effort cocaine self-administration selectively in dependent rats

There was no significant difference between shHCRT or shSCR (*n* = 5, each) rats in ShA cocaine self-administration under an FR1 schedule (Figure 2A; Group: F_(1,9)_= 2.43, p= 0.15.; Session: F_(10,90)_= 0.70, p= 0.72; Group x session: F_(10,90)_= 0.47, p= 0.91). However, upon LgA to cocaine self-administration, shHCRT significantly attenuated cocaine self-administration under both an FR1 schedule (Figure 2B; Group: F_(1,8)_= 6.24, p < 0.05; Session: F_(13,104)_= 1.07, p= 0.39; Group x session: F_(13,104)_= 1.09, p= 0.38) and a PR schedule of reinforcement (Figure 2C; *U*(8) = 0, *Z* = 2.51, p < 0.05). These results indicate that HCRT transmission contributes to cocaine self-administration under low-effort conditions in extended-access rats, but not in limited-access rats.

**Figure 2.**
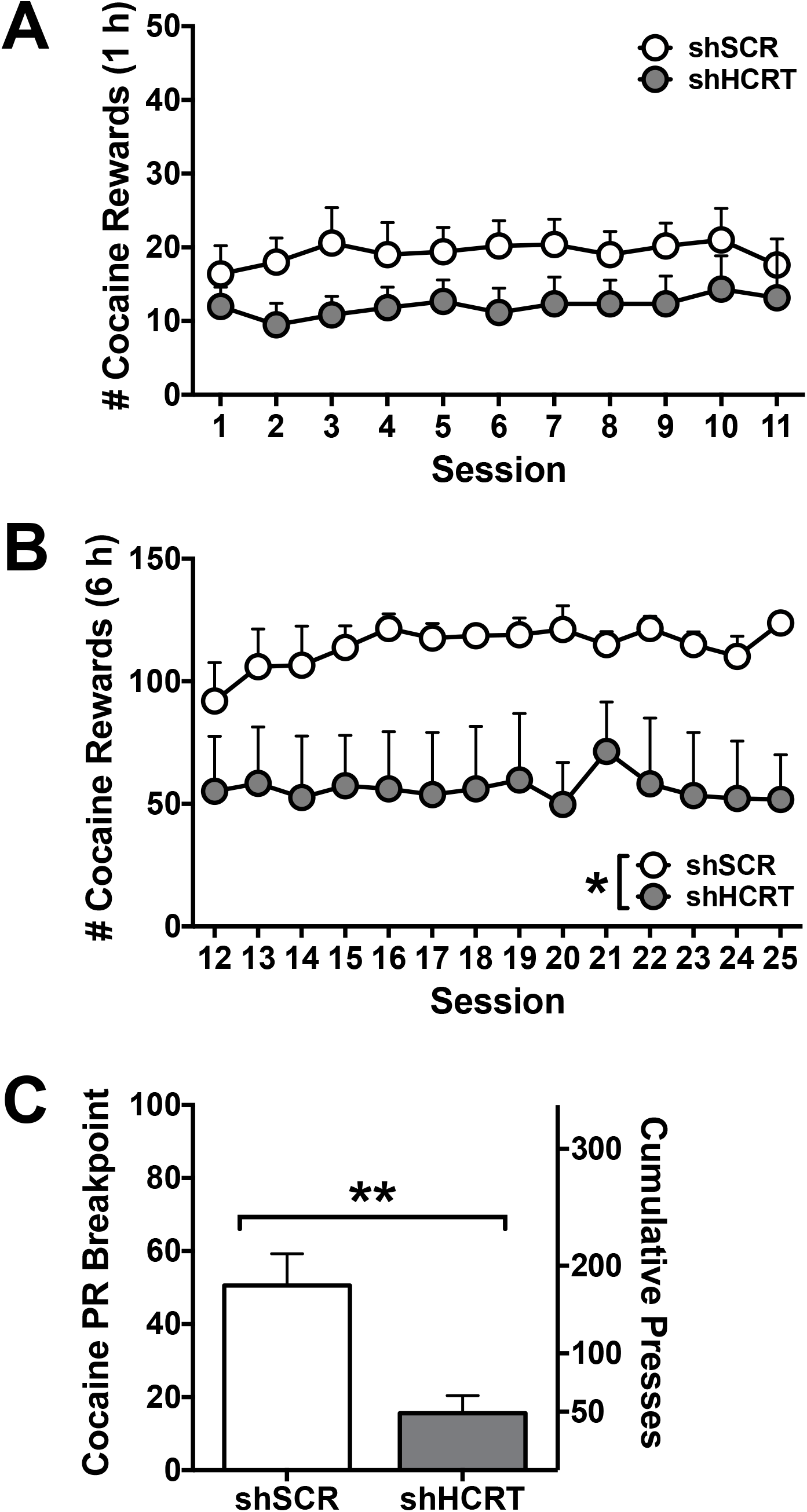
Long-term *Hcrt* knockdown reduces cocaine self-administration in dependent rats. (AB) Symbols represent mean number of cocaine rewards (+SEM) per session. AAV5-shHCRT (*n =* 5) had no effect on cocaine intake over 11 sessions of short access self-administration (A; 1 h; Session 1-11), whereas it significantly reduced cocaine intake over 14 sessions of long access self-administration (B; 6 h; Session 12-25) compared to AAV5-shSCR (*n* = 5). *p<0.05 versus shSCR. (C) Bars represent mean PR breakpoints (+SEM; left axis) and cumulative presses (+SEM; right axis). shHCRT rats showed significantly reduced cocaine intake on a PR schedule of reinforcement compared to shSCR rats. **p<0.01 versus shSCR.

### *Hcrt* knockdown reduces SCM self-administration under PR reinforcement

There was no significant difference between shHCRT or shSCR (*n* = 5, each) rats in SCM self-administration under an FR1 schedule (Figure 3A; Group: F_(1,8)_= 1.07, p= 0.33; Session: F_(11,88)_= 1.71, p= 0.08; Group x session: F_(11,88)_= 0.50, p= 0.90). In contrast, *Hcrt* knockdown significantly attenuated responding for SCM under a PR schedule of reinforcement (Figure 3B; *U*(8) = 2.50, *Z* = 1.98, p < 0.05). These results indicate that the influence of HCRT on the self-administration of palatable food by sated rats is restricted to high-effort conditions.

**Figure 3.**
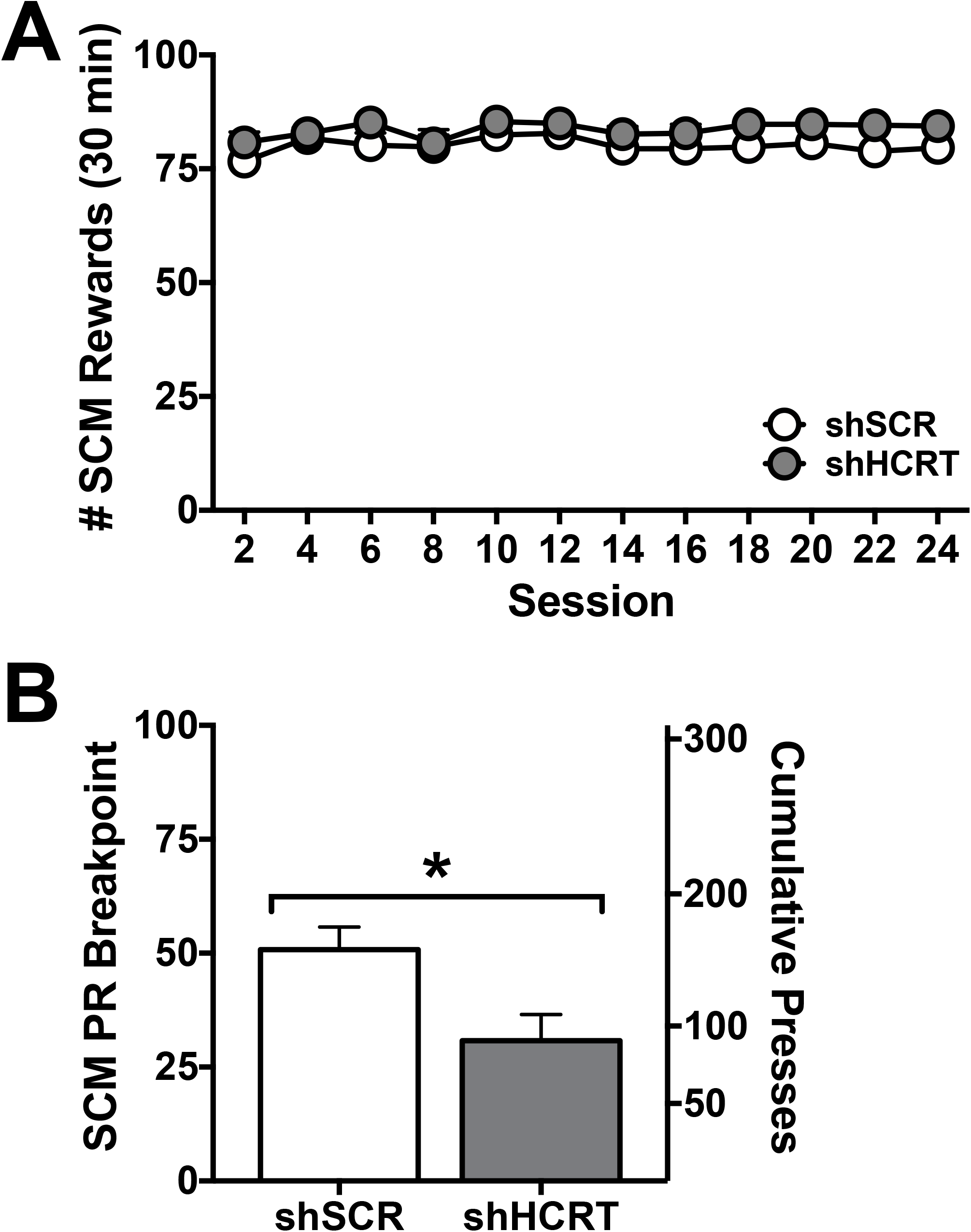
Long-term *Hcrt* knockdown reduces sweetened condensed milk self-administration only under a progressive ratio schedule of reinforcement. (A) Symbols represent mean number of sweetened condensed milk (SCM) rewards (+SEM) per session. AAV5-shHCRT (*n* = 5) had no effect on SCM intake over 24 sessions of self-administration (30 min) compared to AAV5-shSCR (*n* = 5). (B) Bars represent mean PR breakpoints (+SEM; left axis) and cumulative presses (+SEM; right axis). shHCRT rats did show significantly reduced SCM intake on a PR schedule of reinforcement compared to shSCR rats. *p<0.05 versus shSCR.

### *Hcrt* knockdown has no effect on general behavior

There was no significant difference between shHCRT or shSCR (*n* = 5, each) rats in body weight or food pellet self-administration under an FR3 schedule (Figure 4A and B; Body weight: Group: F_(1,8)_= 0.63, p= 0.45; Time: F_(5,40)_= 159.1, p < 0.001; Group x time: F_(5,40)_= 0.46, p= 0.79; Food: Group: F_(1,8)_= 3.48, p= 0.10; Time: F_(1,8)_= 130.50, p < 0.001; Group x time: F_(1,8)_= 0.003, p= 0.96). Similarly, there was no significant difference between shHCRT-and shSCR-treated rats in water self-administration under an FR1 schedule (Figure 4C; Group: F_(1,8)_= 1.16, p= 0.31; Time: F_(1,8)_= 118.20, p < 0.001; Group x time: F_(1,8)_= 1.30, p= 0.29). There was also no significant effect of shHCRT injection on number of crossovers in the activity box compared to shSCR-treated control rats (Figure 4D; Group: F_(1,8)_= 2.20, p= 0.18; Condition: F_(3,24)_= 30.53, p < 0.001; Group x condition: F_(3,24)_= 0.002, p= 0.96) or on the percentage of time spent in the open arms of the elevated plus maze (Figure 4E; *t*(8)= 0.46, p= 0.66). Additionally, shHCRT rats showed no significant difference in latency to thermal nociception on the hot plate test prior to forced-swim stress compared to shSCR rats, and stress produced an equivalent analgesic effect in shHCRT and shSCR rats (Figure 4F; Group: F_(1,8)_= 0.31, p= 0.59; Condition: F_(1,8)_= 86.45, p < 0.001; Group x condition: F_(1,8)_= 1.21, p= 0.30). Finally, there was no significant effect of *Hcrt* knockdown on basal and stress-stimulated corticosterone release (Figure 4G; Group: F_(1,8)_= 0.45, p= 0.52; Condition: F_(1,8)_= 11.97, p < 0.01; Group x condition: F_(1,8)_= 0.92, p= 0.37). Altogether, these data indicate that there is no effect of shHCRT on food/water intake, locomotion, or on measures of anxiety or stress responses in the absence of cocaine. These results demonstrate that the effects of *Hcrt* knockdown on cocaine and SCM self-administration cannot be attributed to a non-specific disruption of behavioral performance or basal stress sensitivity.

**Figure 4.**
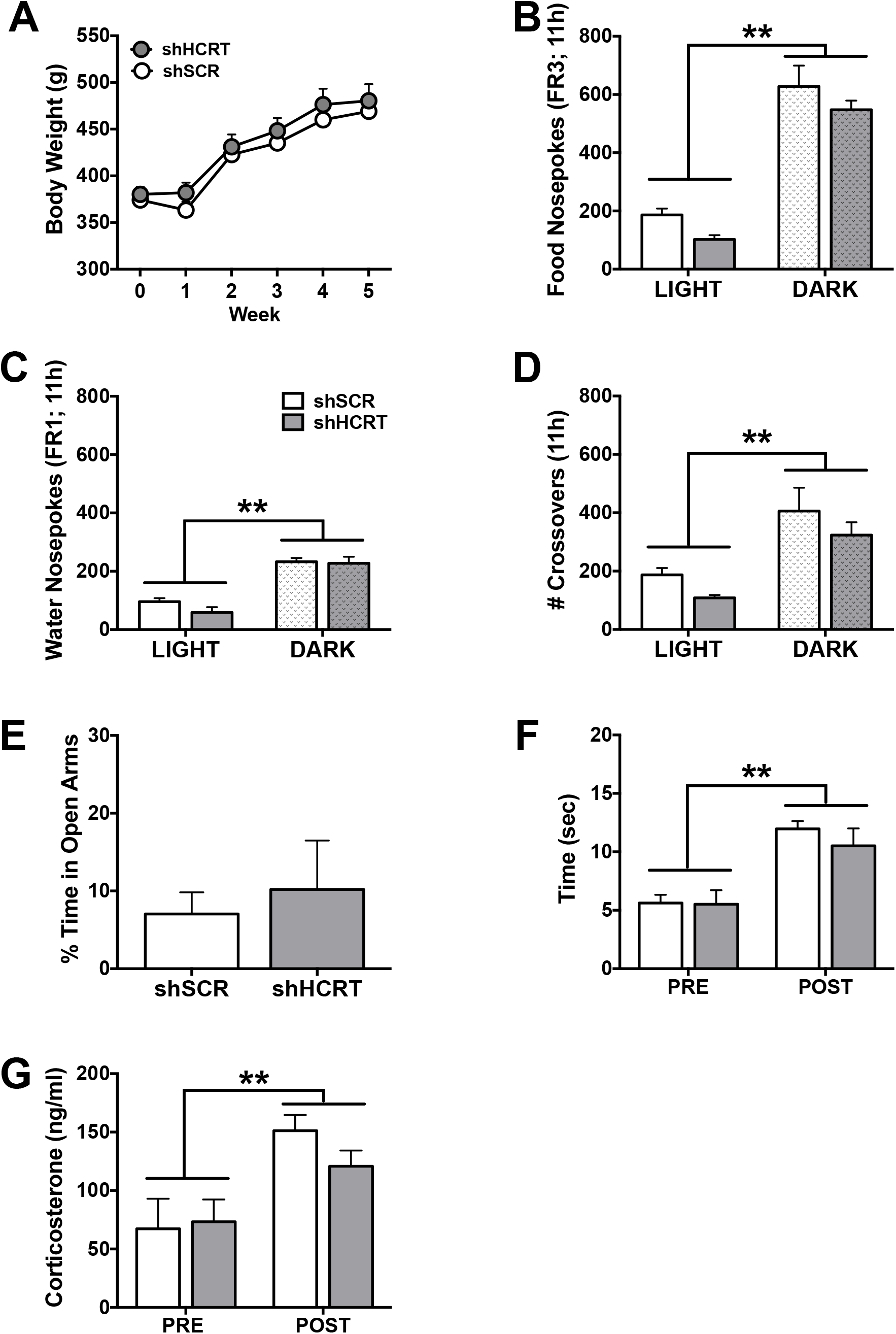
Long-term *Hcrt* knockdown has no effect on general measures of behavior and physiology. (A) Symbols represent mean body weight (g) each week following AAV5-shHCRT or AAV5-shSCR injection (Week 0). There was no significant difference in body weight between shHCRT and shSCR rats (*n* = 5, each). (B-C) Bars represent the mean number of food pellet (B) or water (C) rewards (+SEM) per 11h time epoch. There was no significant difference between shHCRT and shSCR rats in number of food pellets (FR3) or water (FR1) rewards in either the light or dark phase of the circadian cycle. **p<0.01 versus LIGHT. (D) Bars represent the mean number of crossovers (+SEM) in a locomotor activity box per 11h time epoch. There was no significant difference between shHCRT and shSCR rats in number of crossovers in either the light or dark phase. **p<0.01 versus LIGHT. (E) Bars represent percentage of time spent in the open arms (+SEM) of an elevated plus-maze. There was no significant difference in anxiety-like behavior between shHCRT and shSCR rats. (F) Bars represent time (sec; +SEM) to hind paw lick on a 54°C hotplate prior to (PRE) or following (POST) forced swim stressor. There was a significant main effect of stress resulting in an increase of withdrawal latency time following forced swim (stress-induced analgesia), but no significant difference between shHCRT and shSCR rats. **p<0.01 versus PRE. (G) Bars represent mean plasma corticosterone levels (ng/ml; + SEM) prior to (PRE) or following (POST) forced swim stressor. There was a significant effect of stress on corticosterone release, but no significant difference between shHCRT and shSCR rats. **p<0.01 versus PRE.

### Adaptations in local neuropeptide expression

Molecular adaptations of two additional neuronal populations intermingled with HCRT neurons in the dorsal hypothalamus, PDYN and MCH, were analyzed (Figures 5 and 6, respectively). *Pdyn* mRNA expression in the dorsal (i.e., dorsomedial hypothalamus and lateral hypothalamic area) and ventral (i.e., ventromedial hypothalamus and arcuate nucleus) parts of the hypothalamus (Figure 5A) was examined in adjacent sections of brains analyzed for the time-course of *Hcrt* knockdown using the AAV2 serotype. Longterm *Hcrt* knockdown significantly reduced *Pdyn* expression in the dorsal part of the hypothalamus where *Pdyn* is expressed by HCRT neurons 4 weeks following shHCRT injection (Figure 5B; Group: F_(1,5)_= 46.75, p< 0.01.; Time: F_(2,5)_= 3.33, p= 0.12; Group x time: F_(2,5)_= 2.96, p= 0.14), but not in the ventral part of the hypothalamus where there are no HCRT neurons (Figure 5C; Group: F_(1,5)_= 0.82, p= 0.41.; Time: F_(2,5)_= 1.51, p= 0.31; Group x time: F_(2,5)_= 0.15, p= 0.87). MCH-immunoreactivity was examined in adjacent brain sections from the third, general behavior cohort (Figure 6). There was a nonsignificant trend for a reduction in the number of MCH-positive neurons within the dorsal hypothalamus following long-term *Hcrt* knockdown (Figure 6B, *t*_(8)_ = 1.98, p = 0.08).

**Figure 5.**
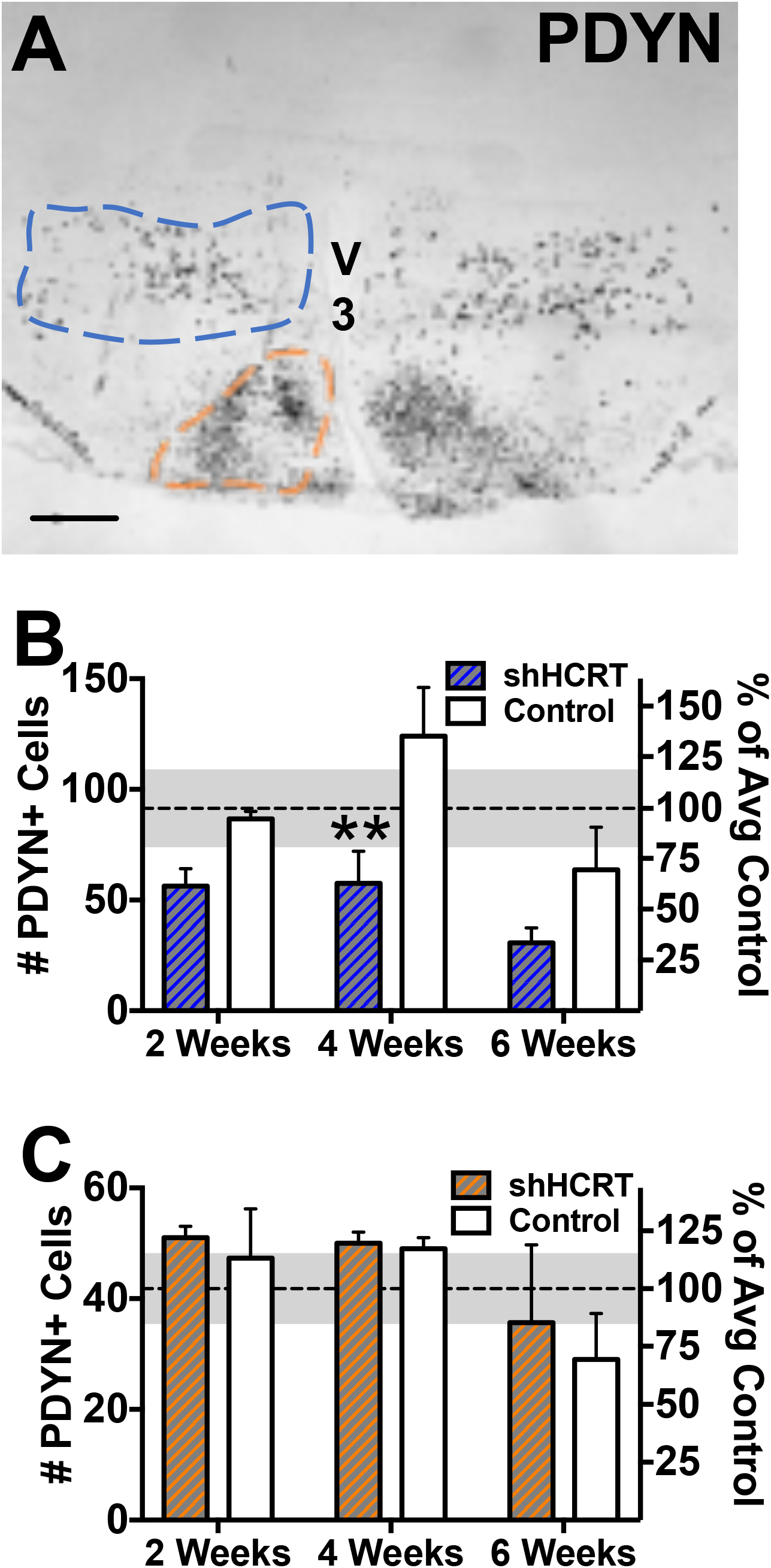
Histological characterization and time-course quantification of *Pdyn* expression following *Hcrt* knockdown. Rats received unilateral double injections of AAV2-shHCRT, with the contralateral hemisphere used as a control site, and brains were collected 2, 4, or 6 weeks later (*n* = 2-3 per time-point). *Pdyn* (A) expression was examined on sections using *in situ* hybridization. V3, third ventricle; scale bar = 500 μm. The number of PDYN+ cells was then quantified within the HCRT neuronal field (i.e., dorsal hypothalamus; blue outline) and on PDYN neurons located more ventrally (orange outline). (B-C) Bars are mean cell counts represented as a percentage of the control hemisphere (% of Control; +SEM) across three coronal sections. (B) HCRT knockdown was accompanied by a concomitant reduction in the number of PDYN-positive neurons in the dorsal hypothalamus (blue outline) at the 4-week time-point. (C) The viral vector had no effect on PDYN neurons located more ventrally (orange outline). Dashed horizontal line is percentage of control averaged over 3 time-points (gray shade ±SEM). **p<0.01 versus contralateral control hemisphere (right).

**Figure 6.**
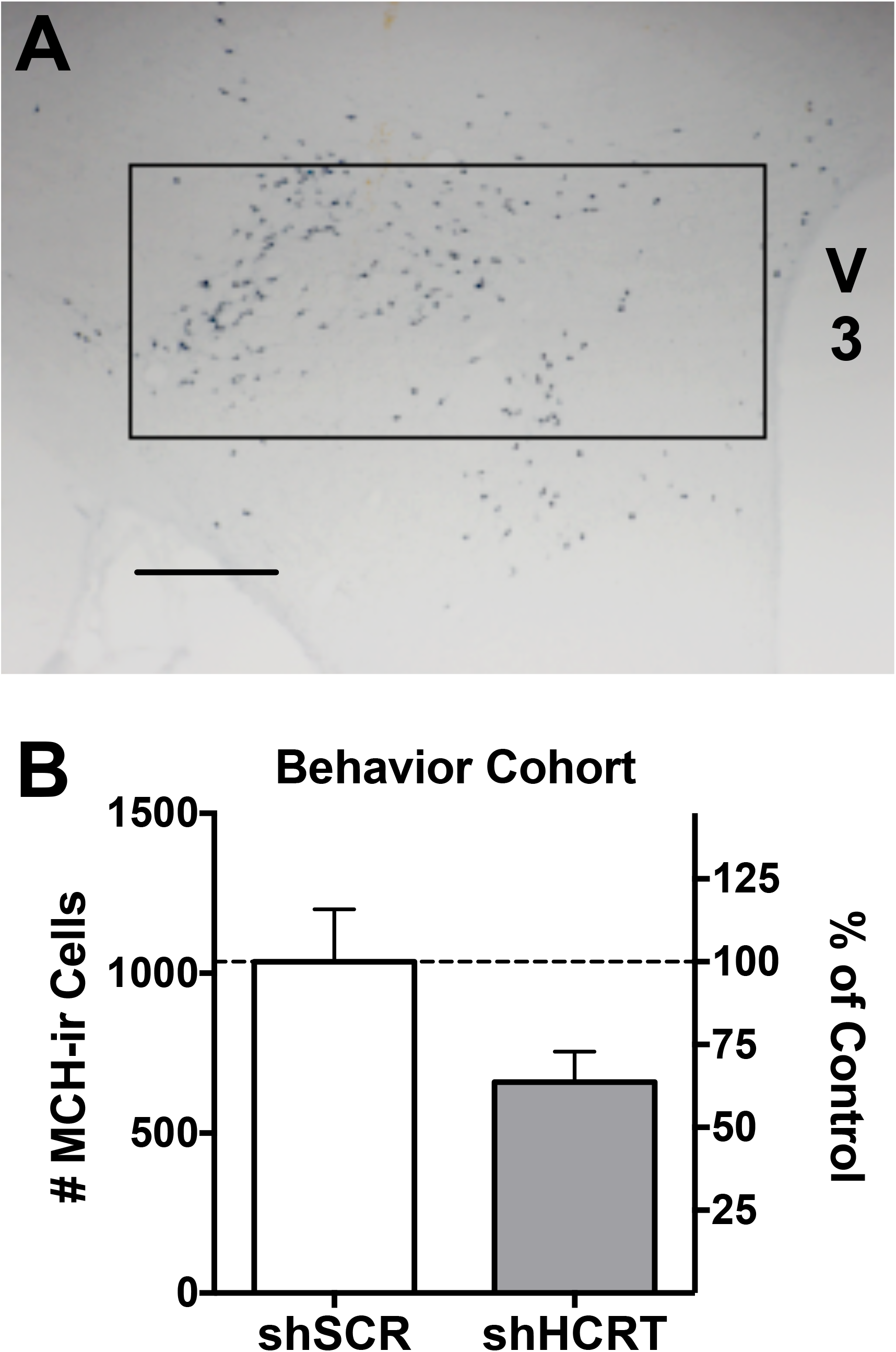
Histological characterization and quantification of MCH cells following *Hcrt* knockdown. Adjacent tissue sections from behavior cohort were processed for MCH-positive immunolabeling. (A) Representative photomicrograph of MCH-immunostaining. V3, third ventricle; scale bar = 500 μm. MCH-immunoreactive (MCH-ir) cells were counted bilaterally within the dorsal hypothamalmus (boxed area) in AAV5-shSCR and AAV5-shHCRT-treated animals (*n* = 5, each). (B) Long-term *Hcrt* knockdown has no effect on MCH-ir cells. Bars represent the mean (+SEM) number of MCH-ir neurons (left axis) or the percentage of the control group (shSCR; right axis) across three coronal sections

## Discussion

The HCRT system has been implicated in the motivation to consume and seek drugs of abuse, but its role in extended-access drug-taking associated with compulsive-like cocaine use is poorly understood. Here, a genetic approach was used to probe the role of HCRT neurotransmission in extended-access cocaine self-administration. Efficient, long-lasting knockdown of *Hcrt* expression throughout the dorsal hypothalamus was achieved by virally-mediated RNA interference in a rat model, circumventing the need to chronically administer HCRT-receptor antagonists. Our results confirm previous pharmacological evidence that HCRT neurotransmission promotes operant responding for both drug and non-drug rewards under motivationally salient, high-effort conditions (Borgland 2009; Mahler *et al*, 2014). Furthermore, the present study demonstrates that HCRT neurotransmission contributes to cocaine intake even under a low-effort contingency when access to cocaine self-administration is extended. Altogether, these data provide novel insights into the role of HCRT in drug use under conditions of pathologic motivation, and validates virally-mediated *Hcrt* silencing as a valuable approach to investigate the behavioral relevance of HCRT neurotransmission in rodent models.

### A role for HCRT in the transition to cocaine addiction-like behavior

Our finding that *Hcrt* knockdown did not affect cocaine self-administration under low-effort conditions (FR1 schedule of reinforcement) in rats offered 1 h daily access to the drug corroborates previous pharmacological studies. HCRT-R1 antagonists show no or minimal effects on low-effort, FR1 responding for cocaine self-administration in animals allowed limited access to cocaine (España *et al*, 2010; Schmeichel *et al*, 2016; Smith *et al*, 2009). Likewise, intracerebral injection of HCRT-1 does not affect FR1 responding for cocaine (Boutrel *et al*, 2005; España *et al*, 2011). **Furthermore, the lack of effect on FR1 responding under short access to cocaine conditions in the current studies is not likely attributed to a lack of *Hcrt* silencing at time of testing (i.e., 2-3 weeks) as we confirmed a nearly 85% knockdown of *Hcrt* as soon as two weeks following AAV injection (see Figure S1).** Altogether, the genetic approach used in the current studies lends further support to the hypothesis that HCRT transmission is not necessary for the primary rewarding effects of the drug under conditions of low incentive-motivation.

In contrast, *Hcrt* silencing strongly reduced FR1 cocaine responding in extended-access rats. This finding suggests, that HCRT neurotransmission may contribute to the activation of stress systems and subsequent dampening of reward systems that are associated with extended access to cocaine self-administration and are hypothesized to drive excessive, compulsive-like cocaine taking in addiction (for review, Koob *et al*, 2014). In support of this, central infusion of HCRT-1 peptide increases thresholds of intracranial self-stimulation, a rate-independent measure of brain reward sensitivity (Boutrel *et al*, 2005; Hata *et al*, 2011), suggesting an inhibitory action on reward systems. Furthermore, central HCRT-1 administration reinstates previously extinguished drug-seeking, whereas HCRT-R1 antagonism readily blocks stress-induced reinstatement of cocaine-seeking (Boutrel *et al*, 2005; Harris *et al*, 2005; Martin-Fardon *et al*, 2010; Matzeu *et al*, 2016; Schmeichel *et al*, 2016; Wang *et al*, 2009; Zhou *et al*, 2012). **Both preclinical and clinical work has established a role for HCRT in stress responses to anxiety-and panic-associated behaviors (Harris *et al*, 2005; Johnson *et al*, 2010, 2012; Suzuki *et al*, 2005). However, whether HCRT-related stress reactivity is sensitized under conditions of chronic cocaine, promoting the hyper-aroused state required for drug-taking, has yet to be** determined. Additionally, it will be interesting to examine the effect of *Hcrt* knockdown on stress-induced reinstatement cocaine-seeking in future studies.

In conclusion, extended-access cocaine self-administration extends the role of HCRT signaling in the motivational activation of cocaine reinforcement, such that HCRT contributes not only to high-effort but also to low-effort responding under conditions where motivation is high. Furthermore, our findings suggest the feasibility of normalizing compulsive cocaine intake in addicted individuals via sustained inhibition of HCRT signaling.

### HCRT in reward-based feeding

*Hcrt* knockdown also decreased the motivation to obtain palatable food (SCM) under a PR schedule of reinforcement, which suggests a role for HCRT in food seeking under high-effort conditions, as well as during the consumption of highly palatable foods. This finding is consistent with the existing literature (for review, Mahler *et al*, 2014). HCRT was initially reported to regulate general feeding behavior by demonstrating that central administration of HCRT-1 and HCRT-2 stimulates feeding (Sakurai *et al*, 1998), whereas blockade of HCRT signaling with an HCRT-R1 antagonist, an anti-HCRT antibody, or genetic inactivation of *Hcrt* reduces food intake (Hara *et al*, 2001; Haynes *et al*, 2000; Yamada *et al*, 2000). Subsequent studies revealed that the modulation of feeding by HCRT is restricted to the light phase of the circadian cycle (España *et al*, 2002; McGregor *et al*, 2011). Our observations are consistent with the latter findings, as *Hcrt* silencing did not significantly affect responding for regular food pellets overall, but there was a trend for reduced food self-administration during the light phase. HCRT is also likely to play an important part in the hedonic aspect of feeding (i.e., feeding beyond caloric needs; Zheng and Berthoud, 2007). Consistent with this hypothesis, central administration of HCRT-1 increases the motivation for palatable food, whereas blockade of HCRT signaling with an HCRT-R1 antagonist attenuates this reward-based feeding (Borgland *et al*, 2009; Choi *et al*, 2010; Thorpe *et al*, 2005). Therefore, together with these studies, our data support the hypothesis that HCRT signaling mediates motivation to obtain palatable food.

### HCRT modulation of arousal-dependent behavior

The present study showed that *Hcrt* knockdown did not affect food or water self-administration, locomotor activity, or sensitivity to tests of anxiety-like behavior and stress-induced analgesia or corticosterone release. At first glance, these results may be surprising considering the established role of HCRT in a multitude of arousal-dependent behaviors and physiological functions including energy homeostasis, sleep/wake state, response to stress, and reinforcer-motivated behavior (for review, see Berridge and España, 2005; Boutrel and de, 2008; Johnson *et al*, 2010; Tsujino and Sakurai, 2009). It is possible that the lack of significant behavioral alterations following virally-mediated *Hcrt* silencing resulted from compensatory activity of the few remaining HCRT-positive neurons, which may have been sufficient to modulate relevant arousal-dependent behaviors. In addition, the fact that *Hcrt* expression was silenced in adult animals may explain the discrepancy with results obtained upon constitutive deletion of *Hcrt* in knockout mice, whose phenotype may partially result from developmental compensations (or review, de Lecea *et al*, 2002).

In the present study, shHCRT-treated rats maintained normal body weights, levels of food and water consumption, and locomotor activity. In particular, there was no detectable effect of *Hcrt* silencing on these measures during the dark phase of the circadian cycle, when cocaine and SCM self-administration sessions were conducted. This rules out performance disruption as a potential explanation for the reduction in operant responding for cocaine and SCM.

In addition, *Hcrt* knockdown did not produce a significant blunting of stress sensitivity in the absence of cocaine, as measured by anxiety-like behavior in the elevated plus maze and thermal analgesia or plasma corticosterone elevation following forced swim stress. This suggests that HCRT neurotransmission does not indiscriminately contribute to all responses to psychological or physiological stressors and that the implication of HCRT signaling in extended-access cocaine self-administration is specific to homeostasis and affective state dysregulation associated with withdrawal from chronic, excessive cocaine intake.

### Molecular neuroadaptations following virally-mediated shRNA interference in the rat

To date, physiological functions of the HCRT neuropeptide system have mainly been investigated using acute pharmacological manipulation or constitutive gene deletion. A limited number of studies have used alternative approaches such as local infusions of small interfering RNAs or antisense morpholinos for the transient knockdown of HCRT (Chen *et al*, 2006; Kim *et al*, 2015; Prasad and McNally, 2014; Reissner *et al*, 2012), or local injection of shRNA-encoding viral vectors for the long-term knockdown of HCRT, HCRT-R1 or HCRT-R2 (Arendt *et al*, 2014; Chen *et al*, 2010, 2013; Choi *et al*, 2012; Sharf *et al*, 2010). Here we report virally-mediated knockdown of HCRT in adult rats. Thus, the novelty of our approach warranted further examination of molecular adaptations potentially elicited by sustained silencing of *Hcrt* expression in the adult brain.

In the dorsal hypothalamus, DYN and HCRT are expressed by the same neurons, being co-released from the same synaptic vesicle (Chou *et al*, 2001; Muschamp *et al*, 2014). Furthermore, these neurons synapse onto each other and local release of DYN tonically inhibits neighboring HCRT neurons (Li and van den Pol, 2006). We found that HCRT knockdown was accompanied by a ~34-53% reduction in the number of *Pdyn*-positive cells in the HCRT neuronal field (i.e., dorsal hypothalamus). Importantly, the shHCRT vector had no effect on *Pdyn* expression in more ventral regions of the hypothalamus, thereby ruling out an off-target effect of the shRNA sequence on *Pdyn* mRNA, or a neurotoxic effect of viral transduction or GFP expression. This observation suggests that *Pdyn* downregulation is a compensatory adaptation that occurs downstream of HCRT knockdown. DYN has been shown to balance the effects of HCRT on neuronal excitability in a subset of ventral tegmental area dopaminergic neurons (Baimel *et al*, 2017; Muschamp *et al*, 2014). Thus, in shHCRT-treated rats, the reduced need to control the activity of HCRT neurons in the absence of excitatory HCRT transmission could have triggered *Pdyn* downregulation. At the behavioral level, HCRT- and DYN-receptor activation both elicit an upward shift of intracranial reward thresholds (Boutrel *et al*, 2005; Todtenkopf *et al*, 2004). It is possible that the downregulation of *Pdyn* associated with HCRT knockdown may have potentiated the amplitude of phenotypic changes observed in shHCRT rats. HCRT-DYN interaction effects on cocaine self-administration is an important topic for future studies.

Additionally, we observed that HCRT knockdown induced a moderate, although not statistically significant, decrease in the number of MCH-positive cells (~36% reduction). MCH neurons are intermingled with HCRT neurons (Broberger *et al*, 1998; Elias *et al*, 1998) and several studies have indicated a direct connectivity between these two cell types, whereby HCRT excites MCH neurons and MCH counters the activation of HCRT neurons (Barson *et al*, 2013; Gao and van den Pol, 2001, 2002; van den Pol *et al*, 2004; Rao *et al*, 2008). The inhibitory tone exerted by MCH on HCRT neurons could explain how MCH downregulation may represent a network adaptation to sustained inhibition of HCRT transmission. A recent study also revealed HCRT neurons inhibit MCH neurons via a local microcircuit involving HCRT-induced GABA release (Apergis-Schoute *et al*, 2015). It is therefore possible that MCH was downregulated to compensate for the loss of HCRT-driven inhibition of MCH neurons in shHCRT rats. At the behavioral level, blocking MCH signaling attenuates cocaine self-administration, as well as cue- and drug-induced reinstatement (Chung *et al*, 2009). Similar effects were reported for alcohol (Cippitelli *et al*, 2010). Thus, in the current studies, MCH downregulation may have contributed to decreasing cocaine intake in shHCRT rats, but this would need to be directly tested in future studies.

Finally, the adaptations elicited by sustained inhibition of HCRT signaling in adulthood may differ from developmental compensations elicited by constitutive gene knockout or genetic ablation of neurons (e.g., Sharf *et al*, 2010). Notably, *Pdyn* expression in the dorsal hypothalamus is unchanged in HCRT knockout mice and MCH neurons are not affected in mice that have lost HCRT neurons shortly after birth (Chou *et al*, 2001; Hara *et al*, 2001). Although counterintuitive, these observations combined with our findings suggest that late-onset downregulation of HCRT signaling is more prone to elicit perturbations in cross-talking neuropeptide systems than the life-long absence of HCRT. An important implication of this interpretation is that chronic administration of a HCRT-receptor antagonist in adult subjects would be expected to produce similar reductions in hypothalamic PDYN and MCH expression, which may in turn enhance the therapeutic efficacy of this approach in the treatment of cocaine use disorder.

## Conclusion

In summary, long-term *Hcrt* silencing via virally-mediated RNA interference yielded novel insights into the role of HCRT transmission in cocaine and palatable food reinforcement without affecting general activity, operant performance, or stress reactivity. *Hcrt* knockdown attenuated cocaine and SCM self-administration selectively under high-effort conditions and reduced cocaine intake under conditions of high-motivation associated with extended-access. These findings suggest a specific functional role for HCRT signaling in compulsive-like cocaine self-administration, as well as in reward-based feeding behavior.

Supplementary information is available at the *Neuropsychopharmacology* website.

## Acknowledgements

**Funding and Disclosures**: This research was supported by grants from the National Institute on Drug Abuse, DA004398 (GFK), DA033344 (RMF), DA036355 (BES), the National Institute of Alcohol Abuse and Alcoholism, AA016658 (BLK), AA024198/AA006420/AA021491 (CC) and AA024146/AA006420 (RMF), and by the Pearson Center for Alcoholism and Addiction Research. A portion of this work was also supported by the Intramural Research Program of the National Institute on Drug Abuse. The authors thank Zhi-Bing You for assistance with corticosterone measurements. The authors declare no conflict of interest. This is article number 29409 from the Scripps Research Institute.

